# A thermodynamic cycle to predict the competitive inhibition outcomes of an evolving enzyme

**DOI:** 10.1101/2025.02.03.636225

**Authors:** Ebru Cetin, Haleh Abdizadeh, Ali Rana Atilgan, Canan Atilgan

**Author notes:** **Correspondence:** Canan Atilgan, Faculty of Natural Sciences and Engineering, Sabanci University, Tuzla 34956 Istanbul, Turkey. Department of Chemistry & Biochemistry, University of Arizona (Ebru Cetin). Department of Strategic Development, University of Twente (Haleh Abdizadeh).

## Abstract

Understanding competitive inhibition at the molecular level is essential for unraveling the dynamics of enzyme-inhibitor interactions and predicting the evolutionary outcomes of resistance mutations. In this study, we present a framework linking competitive inhibition to alchemical free energy perturbation (FEP) calculations, focusing on *E. coli* dihydrofolate reductase (DHFR) and its inhibition by trimethoprim (TMP). Using thermodynamic cycles, we relate experimentally measured binding constants (*K*_*i*_ and *K*_*m*_) to free energy differences associated with wild-type and mutant forms of DHFR with a mean error of 0.9 kcal/mol, providing insights into the molecular underpinnings of TMP resistance. Our findings highlight the importance of local conformational dynamics in competitive inhibition. Mutations in DHFR affect substrate and inhibitor binding affinities differently, influencing the fitness landscape under selective pressure from TMP. Our FEP simulations reveal that resistance mutations stabilize inhibitor-bound or substrate-bound states through specific structural and/or dynamical effects. The interplay of these effects showcases significant epistasis in certain cases. The ability to separately assess substrate and inhibitor binding provides valuable insights, allowing for a more precise interpretation of mutation effects and epistatic interactions. Furthermore, we identify key challenges in FEP simulations, including convergence issues arising from charge-changing mutations and long-range allosteric effects. By integrating computational and experimental data, we provide an effective approach for predicting the functional impact of resistance mutations and their contributions to evolutionary fitness landscapes. These insights pave the way for constructing robust mutational scanning protocols and designing more effective therapeutic strategies against resistant bacterial strains.

## Introduction

Predicting competitive inhibition fates in point mutants of proteins is critical for the process of drug discovery whereby one strives for inhibitors that are effective on a wide range of variants.^1^ This strategy would be critical for developing drugs which have efficacy within the time frames when resistance mutations arise and get permanently fixed.^2^ While accurate prediction of binding affinities helps prioritize compounds with the highest likelihood of success, reducing trial-and-error in drug discovery, experimental determination of binding affinities and development of suitable biochemical assays are expensive and time-intensive.^3^ Paradoxically, high failure rates in clinical trials often stem from a poor understanding of binding affinities or off-target effects. One route to optimize these efforts is to develop computational approaches that allow virtual screening of compounds, identifying promising candidates early and reducing the costs of wet-lab experiments.

The level of accuracy and precision required for such predictions are currently provided by free energy perturbation (FEP) calculations.^4^ While accuracies below 0.5 kcal/mol are desired for reliable predictions, current level of confidence provided by FEP calculations is around 1 kcal/mol.^5^ Improved accuracy over traditional methods such as docking or molecular mechanics force fields which make FEP more reliable in early-stage drug design stems from its full atomistic resolution which enables modeling complex thermodynamic contributions like solvation and entropic effects.^6^ Conversely, simpler models often take such critical factors into consideration only as mean field effects, sacrificing from accuracy and precision. FEP-based approaches are also adept at capturing small changes in molecular structure such as substituent modifications and their impact on binding affinity. This feature is critical in lead optimization, where small structural modifications to drug candidates influence activity.

While FEP calculations are costly compared to lower resolution approaches, advancements in molecular dynamics (MD) simulations and GPU computing as well as fine tuning of existing methods have made them ever more scalable and accessible.^7, 8^ The ongoing development of FEP-based methods further ensures that these techniques keep pace with the increasing complexity of drug targets, allowing integration of large-scale FEP applications to pharmaceutical pipelines addressing real life problems.^9^

Despite the many advances, constructing the thermodynamic cycles which are suitable to make decisions on drug efficacy facing various mutations on the target is far from straightforward. Compared to transport or structural proteins, enzymes pose additional challenges due to the kinetic effects that need to be accounted for during lead optimization. Here, we relate binding free energy differences to experimentally determined biochemical constants *K*_*m*_ and *K*_*i*_ measured under the evolutionary pressure of competitive inhibitors. We demonstrate the utility of this approach on the mutants of *E. coli* dihydrofolate reductase (DHFR) (Figure 1a) which catalyzes dihydrofolate (DHF) to tetrahydrofolate (THF) (Figure 1b). We have chosen *E. coli* DHFR due to the extent of the evolutionary trajectories accumulated on this model system under the pressure from the competitor inhibitor trimethoprim (TMP) and the accompanying biochemical data for the emerging mutants.^10, 11^ Moreover, the analyses of precatalytic conformers of DHFR mutants have enabled us and others develop leads that steer the evolutionary trajectories away from the mutational pathways with extreme resistance to the drug^12-14^ making it a testbed for developing drug optimization methodologies. We show that the thermodynamical cycle we propose is a good predictor of mutant fates in competitive inhibition and opens the way forward for high-throughput applications^15^ in predicting mutants arising under various evolutionary pressure scenarios.

**Figure 1.**
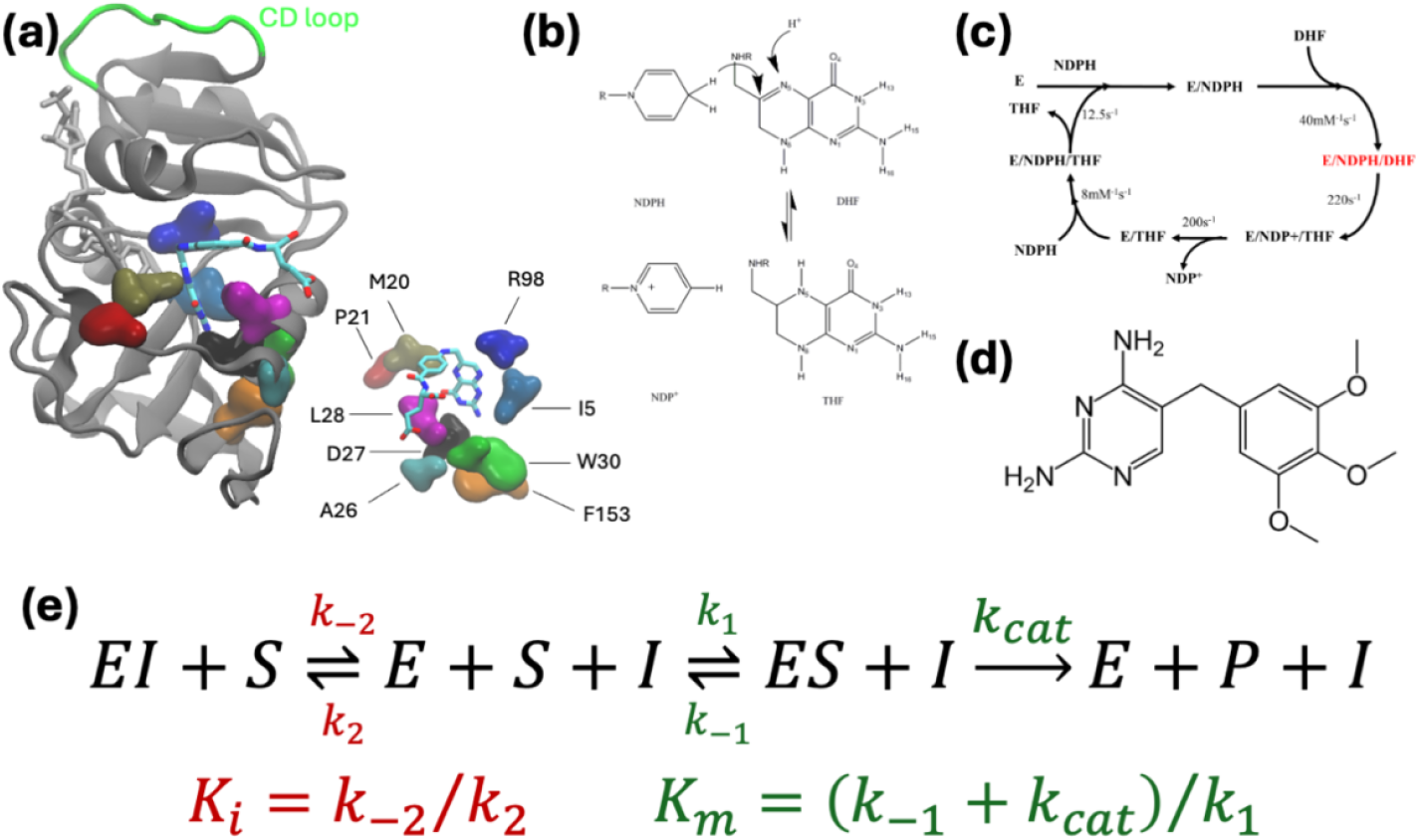
**(a)** DHFR structure displaying the cofactor NDPH (licorice representation colored gold). The substrate DHF and the residues most frequently observed in mutants arising in the morbidostat are also shown from a different perspective on the lower right. DHF is colored by atom type, the mutating residues are shown as volumetric blobs. CD loop is marked in green; **(b)** hydride transfer step; **(c)** catalytic cycle for the enzyme; **(d)** competitive inhibitor studied in this work, trimethoprim (TMP); **(e)** reaction scheme for competitive inhibition; *K*_*i*_ is the inhibitor’s dissociation constant, and *K*_*m*_ is the Michaelis constant.

## Methods

### System Preparation

All simulations for the systems are conducted with the NAMD program.^16^ The initial systems are based on the 1RX2 PDB coded crystal structure^17^ where the cofactor NADPH and folate are bound to the enzyme. The TMP-bound systems are obtained by removing folate and docking TMP to the binding site. For both the DHF-bound and TMP-bound systems, each mutation is introduced with the VMD Mutator Plugin.^18^ Both Charmm22 with CMAP corrections and Charmm36 parameter sets for proteins are utilized.^19^ We report full results from the former in this manuscript, but we verified that we have the same energy differences for the TMP bound M20I, A26T, D27E and I94L systems within error bars. DHF is simulated in a protonated form that models the precatalytic state; thus, the 5-protonated 7,8-dihydrofolate force field parameters were used as reported in the literature.^20^

TMP is modeled in the protonated state as established in our previous work where its force-fields parameters are also listed.^21^ The water box is set to the dimensions of 65×87×65 Å with a padding of at least 10 Å TIP3P water layer in each direction of the protein. The salt concentration is set to isotonic conditions, 0.15 M, with K^+^ and Cl^+^ ions. Particle mesh Ewald summation is utilized to calculate long-range electrostatics with a cutoff distance of 12 Å and a switching distance of 10 Å. The RATTLE algorithm is applied to constrain bonds, and the Verlet algorithm is used with a time step of 2 fs. The temperature is controlled at 310 K by Langevin dynamics with a dampening coefficient of 5 ps^−1^. The pressure is set to 1 atm and regulated by the Langevin piston.

To get initial coordinates for FEP simulations, we conduct classical MD simulations on DHF-bound and TMP-bound WT systems. Due to convergence issues (see Results), equilibration simulations for the W30R and F153S mutants are also carried out. These systems are minimized for 10,000 steps. The resulting structures are subjected to 200 ns long production runs in the *NPT* ensemble which we have shown in our previous work to be ample to equilibrate the mutants.^21, 22^ In another study on DHFR, the last 2 ns of 10 ns long MD runs have been extracted to compute properties of the system.^14^ We therefore utilize various structures at 10 ns time point and beyond, extracted from our 200 ns trajectories, as initial structures for the FEP simulations.

### Alchemical free energy perturbation calculations

The alchemical free energy perturbation method with Zwanzig’s formulation is followed for all the systems listed in Table 1.^23^ The implementation in the NAMD suite of programs is used.^24^ The best practices outlined by Mey *et al*.^25^ are applied. Force field parameters and simulation conditions are the same as those in the previous subsection.

**Table 1.** Mutations for which FEP calculations are carried out.

| System | No. of runs for $\Delta G_s$ | No. of runs for $\Delta G_i$ | Notes |
| --- | --- | --- | --- |
| I5F | 7 | 7 | – |
| M20I | 4 | 4 | – |
| A26T | 4 | 4 | – |
| D27E | 4 | 7 | – |
| L28R<sup>†</sup> | 6/6 | 4/4 | – |
| W30G | 5 | 7 | – |
| W30R<sup>†</sup> | 14/11 | 10/10 | Merge W30R & R30W simulations |
| I94L | 4 | 4 | – |
| F153S | 6 | 6 | S153F simulations used for $\Delta G_s$ |
| L28R-A26T | 7 | 7 | A26T mutation added to L28R |
| L28R-W30G | 7 | 7 | W30G mutation added to L28R |
| W30G-A26T | 6 | 7 | A26T mutation added to W30G |
| I94L-A26T | 7 | 7 | A26T mutation added to I94L |

<sup>†</sup> charge annihilation/creation runs are conducted separately and merged according to <sup>26</sup>.

Systems selected from various equilibration MD simulation time points which are at least 10 ns apart are minimized followed by 0.5 ns equilibration. Wild type (WT) to mutant, mutant to WT conversions, phase-space overlap versus sampling time, the number of *λ* windows, and the effect of equilibration in each *λ* windows were inspected. For the system at hand, we found *λ* = 32 windows to yield consistent results which we present in this work. Each window size is set to 200 ps, with the initial 50 ps being discarded for equilibration. Hence the production in each window is 150 ps long. The systems are run both in forward and backward directions.

To reduce singularities occurring during Leonard-Jones potential calculations, a soft-core potential is introduced at the middle *λ*. The charge changing mutations are problematic because the results depend on box size due to the long-range effects of electrostatic interactions. The protocol proposed by Morgan and Massi^26^ is used for charge correction whereby one set of charge creation and another set of charge annihilation FEP runs are carried out. Since the two charge changing mutations in this work are from a neutral side chain to a positively charged one (L28R and W30R) the created/annihilated charge is a chloride ion. The arithmetic average of the values obtained from these simulations provides the expected free energy difference for the mutation, free from box size effects, while the difference provides the free energy cost of charge creation in the water environment, Cl^-^ in this case. The convergence of the latter value across FEP runs provides an additional check for the convergence of the free energy cost of the mutation.

Different initial structures are used in sampling to enhance the coverage of the potentially available conformational states. At least four different poses are chosen from the classical MD trajectory. These are from the 10, 50, 70, 100, 110, 150, 170 and 200 ns time points obtained from the trajectories described in the previous subsection and whose forward and backward simulations are completed in a stable manner. If additional sampling is necessary due to insufficient convergence of the free energy window overlaps, more simulations selected from these time points are added to the pool. The acquired data are merged and analyzed by Bennett-acceptance ratio (BAR) method ^27^ as implemented in the alchemlyb library.^28^ Errors are calculated from a simple RMSD averaging of the errors of the individual BAR calculations. All outputs and codes used in their analyses are provided on Github (see data availability statement for details).

## Results

### Relating competitive inhibition to alchemical free energy calculations

The shifts in the conformational dynamics due to changes in substrate-enzyme interactions has recently been scrutinized for β-lactamase and underscores the effect of local dynamics on epistatic outcomes.^29^ It is therefore crucial to construct the thermodynamics cycles suitable for the problem at hand whereby the reaction step(s) effective on local conformational dynamics be included in the calculations while those that are independent of mutation types be assumed constant over the systems.

We will focus on the competitive inhibition of *E. coli* DHFR (Figure 1) by the inhibitor TMP. Ample biochemical experimental data have been published for this enzyme.^10^ DHFR converts dihydrofolate (DHF) to tetrahydrofolate (THF) by the transfer of one proton from the cofactor NDPH (Figure 1b) and another from the solvent environment. The catalytic cycle of *E. coli* DHFR consists of five steps (Figure 1c): Starting with the initial binding of the cofactor NADPH, binding of DHF to the NADPH-enzyme complex to form a ternary complex follows. A hydride ion is then transferred from NADPH to DHF, reducing it to tetrahydrofolate (THF) and oxidizing NADPH to NADP^+^. Once the product THF is released from the enzyme, NADP^+^ dissociates from the enzyme, allowing it to bind new substrates. The hydride transfer step is rapid and not rate-limiting. Instead, the release of NADP^+^ is slower and often determines the overall rate of the catalytic cycle.^30^ The Met20 loop undergoes significant conformational shifts during the catalytic cycle.^17^ After hydride transfer, the loop must reopen to allow NADP^+^ to exit the active site. This conformational change is energetically demanding and slows down the release of NADP^+^. In the precatalytic step, alignment for hydride transfer occurs. Precise positioning of NADPH and DHF allows for optimal orbital overlap necessary for the hydride transfer from NADPH to DHF. The enzyme’s conformational adjustments lower the activation energy by stabilizing the transition state, making the hydride transfer more efficient.^31-33^

Competitive inhibitors such as TMP (Figure 1d) target DHFR by mimicking the interactions of DHF at the active site with much higher affinity (nM *vs*. μM). Mutations affecting the precatalytic steps can alter binding affinities and conformational dynamics, leading to decreased inhibitor efficacy and drug resistance.^10, 21^ The importance of the precatalytic step in DHFR underscores the intricate interplay between enzyme structure, dynamics, and function. By studying these early events in the catalytic cycle, we can better comprehend how enzymes achieve specificity and efficiency, and how these processes can be manipulated for therapeutic purposes.

The reaction scheme for competitive inhibition is given in Figure 1e, with *K*_*m*_ being the Michaelis constant, and *K*_*i*_ the inhibitor’s dissociation constant. Note that the latter is a purely thermodynamic quantity, whereas the former is kinetic since the product rate constant, *k*_*cat*_, contributes to this quantity. We define the free energy difference relating the relative *K*_*m*_/*K*_*i*_ values of the WT and the mutant protein, ΔΔ*G*_*competition*_,

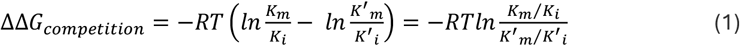

for assessing the relative efficacy of the inhibitor in competitive inhibition. The ratio *K*_*m*_/*K*_*i*_ reflects how much more likely an enzyme is to bind to the substrate than the inhibitor, aiding in understanding the potency of the inhibitor in biochemical contexts. Similarly, the ratio *K*^′^_*m*_/*K*^′^_*i*_ displays the binding propensity of a mutated enzyme toward the substrate relative to the inhibitor. The difference between these ratios quantifies the energetic advantage conferred by the mutation compared to the WT enzyme. In Table 2, we present the *K*_*m*_ and *K*_*i*_ data for *E. coli* DHFR for the WT enzyme and for those mutants that emerge when the bacteria are under evolutionary pressure from TMP. The nine single mutants that appear most frequently in morbidostat experiments are studied in this work;^10^ only P21L and R98P are left out, since they involve changes from/to a proline residue for which FEP simulations are currently not feasible due to the ring structure fused with the protein backbone. We also include four of their double mutant combinations for which biochemical data are available,^10^ leading to a test set of 13 mutants.

**Table 2.** Biochemical data for the mutants studied.

| Mutants | $K_m$ (μM)<sup>†</sup> | $K_i$ (nM)<sup>†</sup> | $\Delta\Delta G$ (kcal/mol)<sup>‡</sup> |
| --- | --- | --- | --- |
| **WT** | **2.86** | **4.59** | -- |
| I5F | 7.68 | 11.90 | 0.02 |
| M20I | 3.57 | 2.90 | 0.42 |
| A26T | 7.65 | 10.10 | 0.12 |
| D27E | 56.40 | 20.21 | 0.92 |
| L28R | 0.95 | 61.94 | -2.28 |
| W30G | 9.49 | 10.20 | 0.25 |
| W30R | 4.97 | 10.53 | -0.17 |
| I94L | 14.87 | 19.15 | 0.14 |
| F153S | 11.32 | 17.00 | 0.04 |
| L28R-A26T | 1.87 | 309.07 | -2.86 |
| L28R-W30G | 0.65 | 309.00 | -3.51 |
| W30G-A26T | 14.87 | 96.77 | -0.86 |
| I94L-A26T | 11.05 | 88.44 | -0.99 |

<sup>†</sup> data reported in ref. <sup>10</sup>
<sup>‡</sup> calculated via eq. 1

Table 2 underscores the fact that, while antibiotic resistance through target modifications is often associated with decreased drug and substrate affinities caused by mutations, experimental measurements suggest the presence of additional resistance mechanisms. Moreover, *K*_*i*_ values alone are insufficient to fully explain TMP resistance. Within the bacterial cell, various other factors including DHFR abundance, catalytic efficiency, thermal stability, nutrient and metabolite availability, excess DHF accumulation, and the demand for THF play significant roles in determining bacterial fitness in the presence of TMP. Of the mutants listed in Table 2, those involving L28R and W30R are the most frequently observed in the morbidostat as the first replacement in the coding region,^10, 22^ consistent with the listed ΔΔ*G*_*competition*_ values calculated by applying equation 1. Moreover, multiple mutants containing the L28R mutation are amongst the toughest to eradicate by drugs,^13^ confirming the further lowered values as listed in Table 2.

We next seek a thermodynamic cycle that will be useful for the prediction of the quantity in equation 1. Under the steady state assumption which is valid when product formation is linear in time, we arrive at the relation for the concentration of inhibitor bound enzyme,

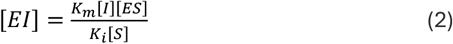

Our aim is to use classical force fields for an enzyme to predict the relative effect of point mutations to the evolutionary outcomes of its overall function. We therefore focus on the equilibrium between the inhibitor bound (enzyme *EI*) and substrate bound enzyme (*ES*). Writing this equilibrium situation twice, once for the WT enzyme (*E*) and once for the mutant (*E*′) we arrive at the thermodynamic cycle depicted in Figure 2. The horizontal equilibria are for the competition of the substrate and the inhibitor for the same binding site in the enzyme *E*, Δ*G*_*E*_, and the mutant enzyme *E*′, Δ*G*_*E*′_. The vertical equilibria represent the cost of mutation in the inhibitor bound enzyme, Δ*G*_*I*_, and the substrate bound enzyme in the precatalytic step, Δ*G*_*s*_. It is these vertical values that may be obtained via FEP simulations.

**Figure 2.**
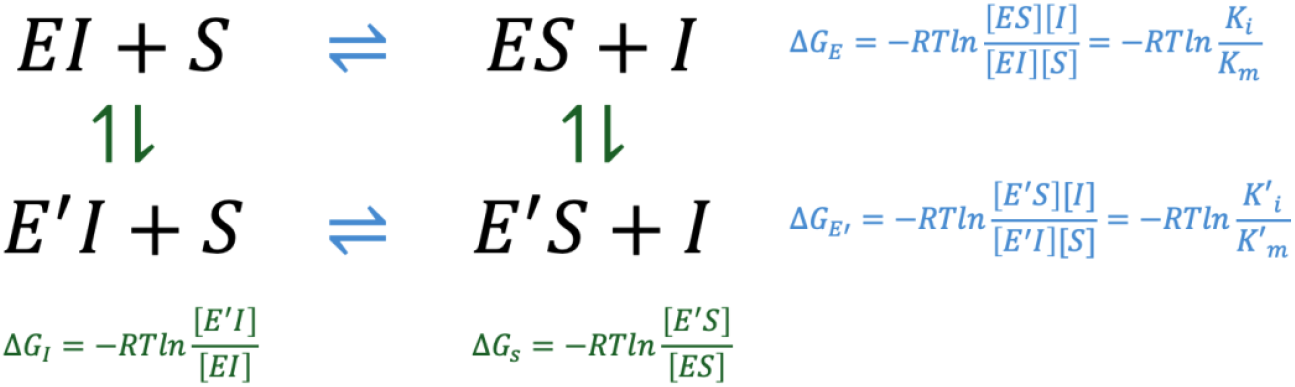
Thermodynamic scheme depicting a cycle for which the alchemical FEP simulations are feasible for the vertical (green) reactions while the substrate – inhibitor exchange reactions occurring in physical reality are shown in the horizontal (blue) reactions. Note that the top horizonal reaction is the 1^st^ and 3^rd^ steps of the competitive inhibition scheme in Figure 1e. The last equalities in the blue equations follow from equation 1.

The sum over the free energy differences around the cycle is zero:

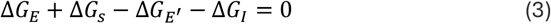

Rearranging and substituting equation 2,

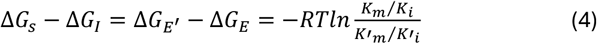

Equation 4 is our final expression linking the FEP simulations to biochemical experimental results which we defined in equation 1. We use FEP simulations to introduce a point mutation to the WT enzyme. We carry out the FEP simulations on both the substrate bound (*ES*) and the inhibitor bound (*EI*) form of the enzyme. The difference in the free energies is expected to provide the right-hand-side, which is obtained from experimentally measured quantities. Our main assumption is that the mutations make a difference predominantly at the precatalytic step when the enzyme samples conformations that are best suited for the catalysis to take place,^34-36^ while the product release is not dependent on the mutations on the enzyme but enters the final observed reaction rates as a common multiple in all mutants bridging the time scale difference.^37^

### DHFR mutation fates predicted by FEP simulations

The calculated values for our case study of competitive inhibition of *E. coli* DHFR by TMP are listed in Table 3. These are visualized against the experimental counterpart values from Table 2, in Figure 3. Our findings point to the applicability of our approach with high accuracy and further disclose the mechanisms that lead to these fates.

**Table 3.** Calculated FEP values.

| System | $\Delta G_s$ (kcal/mol) | $\Delta G_i$ (kcal/mol) | $\Delta\Delta G$ (kcal/mol)<sup>†</sup> |
| --- | --- | --- | --- |
| I5F | -0.66 ± 0.08 | -1.59 ± 0.08 | 0.93 ± 0.11 |
| M20I | 7.94 ± 0.07 | 7.42 ± 0.07 | 0.52 ± 0.10 |
| A26T | -1.90 ± 0.07 | -1.28 ± 0.07 | -0.62 ± 0.10 |
| D27E | 10.02 ± 0.16 | 9.91 ± 0.18 | 0.12 ± 0.24 |
| L28R | -45.62 ± 0.62 | -43.40 ± 0.26 | -2.22 ± 0.36 |
| W30G | 4.96 ± 0.15 | 4.79 ± 0.11 | 0.17 ± 0.19 |
| W30R | -38.81 ± 0.21 | -38.09 ± 0.20 | -0.73 ± 0.29 |
| I94L | -3.97 ± 0.06 | -3.86 ± 0.07 | -0.11 ± 0.09 |
| F153S | -4.22 ± 0.09 | -2.40 ± 0.09 | -1.82 ± 0.13 |
| L28R-A26T | -48.27 ± 0.26 | -43.94 ± 0.26 | -4.33 ± 0.37 |
| L28R-W30G | -41.68 ± 0.28 | -38.34 ± 0.28 | -3.35 ± 0.40 |
| W30G-A26T | 0.15 ± 0.16 | -0.28 ± 0.12 | 0.43 ± 0.20 |
| I94L-A26T | -5.76 ± 0.08 | -5.58 ± 0.09 | -0.18 ± 0.12 |

<sup>†</sup> calculated via eq. 4**Table 4.** Role of epistasis for individual binding events of substrate and inhibitor.<sup>†</sup>

**Figure 3.**
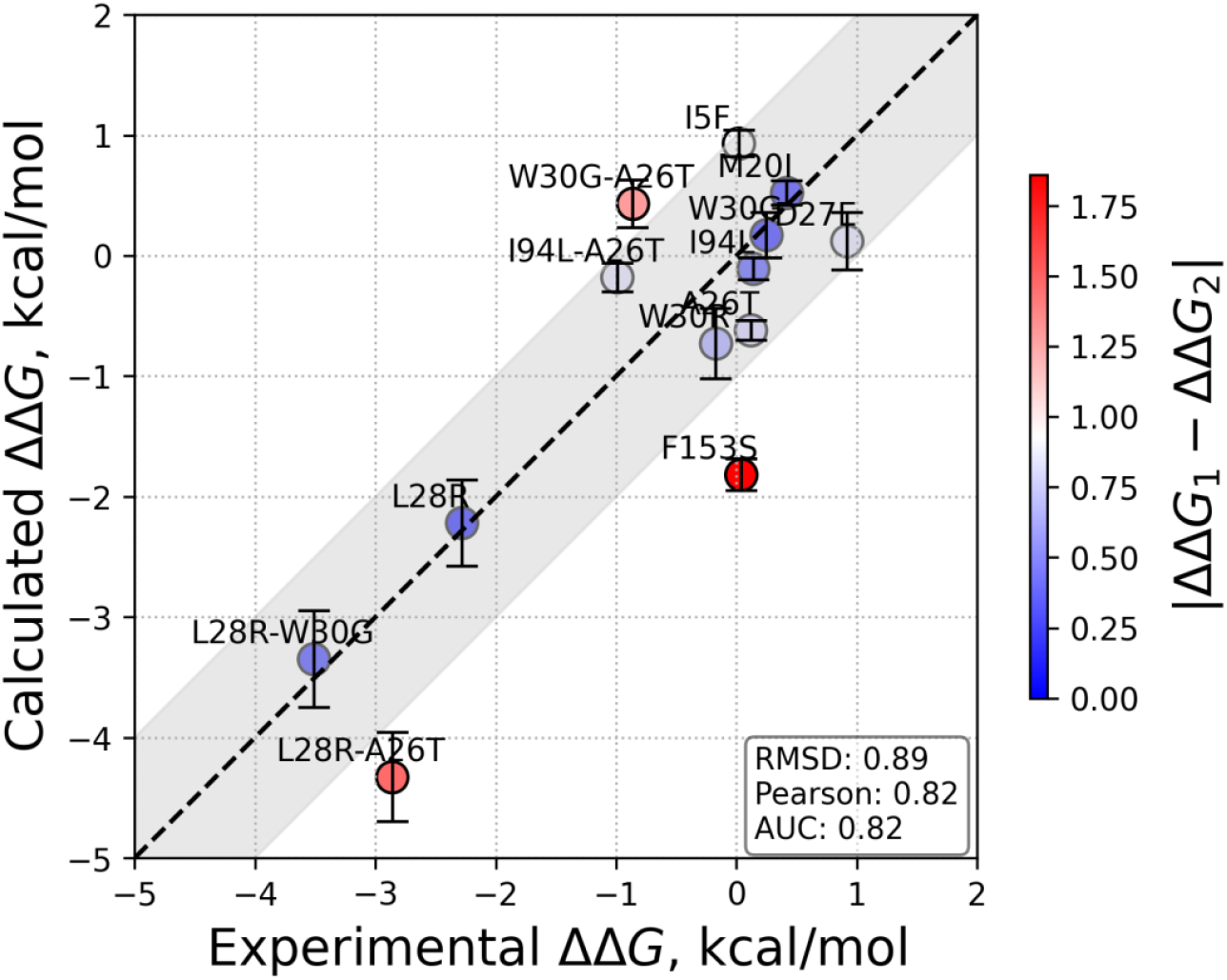
Calculated vs. experimental average ΔΔ*G* values from Tables 2 and 3 with the ±1 kcal/mol error margin shown between the dashed lines. The measures of deviations in terms of RMSD of the data, Pearson correlation coefficient and AUC are provided in the inset.

Inspection of the experimental data in Table 2 shows that according to equation 1, there is only one mutation that displays significant stabilization of the mutant while in competition with TMP. This is L28R which has ΔΔ*G*_*competition*_ = -2.28 kcal/mol. The double mutants L28R-A26T and L28R-W30G are further stabilized, along with W30G-A26T and I94L-A26T which are slightly stabilized by ca. 1 kcal/mol. We find that the mutants destabilize the inhibitor in all but one case (M20I), as one would expect in a resistance mutation, i.e. they display larger *K*_*i*_ than the WT. However, in all single mutants except L28R, the substrate bound complex *ES* is also destabilized with respect to the WT. The latter is an expected outcome, since the competitive inhibitor targets the same modified binding site. The net effect is a ΔΔ*G*_*competition*_ value within 1 kcal/mol of the WT; and this may be the main reason why these usually appear as second or later mutations in morbidostat trajectories. The surviving mutants are then those losing less binding affinity to the substrate than to the inhibitor.

One may indeed follow this outcome from the FEP results with a protein-centric perspective rather than the binder-centric perspective of the experiments. The cost of a mutation may be positive or negative in the presence of the substrate or the inhibitor, as shown by the range of values taken on by Δ*G*_*S*_ and Δ*G*_*I*_ (Table 3). The charge introducing mutations are stabilizing for the protein in the presence of both DHF and TMP (e.g. L28R and W30R), while charge maintaining ones may be positive or negative, depending on the local environment and how the dynamics are altered. Nevertheless, the effect of the mutation is always in the same direction for the DHF and TMP bound forms, and the final stabilization is decided upon by the relative effect of these two values.

### Interpreting epistasis for individual binders and the overall outcomes

Mutational robustness refers to the ability of a genotype to maintain fitness despite mutations, by creating neutral networks in the fitness landscape where multiple genotypes have similar fitness. There is ongoing discussion in the literature regarding the extent to which protein sequence-function relationships are governed by pairwise interactions and higher-order effects.^38, 39^ The answer to this question is important because it determines the ease with which evolutionary landscapes may be constructed, both experimentally and computationally. Interpreting fitness landscapes requires integrating molecular-level interactions, mutation effects, and broader evolutionary dynamics. While free energy differences due to changes in atomic level interactions are far from being the only contributors to the fitness values, they are certainly amongst the most important.^40^ Moreover, dissecting the contributions from the individual sources is also required which we do in Table 4 for the double mutants studied in this work.

**Table 4.** Role of epistasis for individual binding events of substrate and inhibitor.^†^.

| Mutants | $\Delta G_s$<br>(kcal/mol) | $\Delta G_s^e$<br>(kcal/mol) | epistasis in<br>substrate<br>bound DHFR | $\Delta G_i$<br>(kcal/mol) | $\Delta G_i^e$<br>(kcal/mol) | epistasis in<br>inhibitor<br>bound DHFR |
| --- | --- | --- | --- | --- | --- | --- |
| L28R-A26T | -48.27 ± 0.26 | -47.52 ± 0.62 | -0.75 ± 0.68 | -43.94 ± 0.26 | -44.68 ± 0.27 | 0.74 ± 0.37 |
| L28R-W30G | -41.68 ± 0.28 | -40.66 ± 0.64 | -1.02 ± 0.70 | -38.34 ± 0.28 | -38.61 ± 0.28 | 0.27 ± 0.40 |
| **W30G-A26T** | 0.15 ± 0.16 | 3.06 ± 0.17 | **-2.91 ± 0.23** | -0.28 ± 0.12 | 3.51 ± 0.13 | **-3.79 ± 0.18** |
| I94L-A26T | -5.76 ± 0.08 | -5.87 ± 0.09 | 0.11 ± 0.12 | -5.58 ± 0.09 | -5.14 ± 0.10 | -0.44 ± 0.13 |

<sup>†</sup> The superscript e indicates the expected free energy difference if the individual mutations were additive.**Table 5.** Role of epistasis in competitive inhibition.<sup>†</sup>

We compare the calculated versus expected Δ*G*_*S*_ and Δ*G*_*I*_ values, the latter by simply adding the individual free energy differences of the single mutants listed in Table 3. We find that the W30G-A26T double mutation is highly epistatic both for DHF and for TMP binding. Looking at individual contributions, the W30G mutation makes a positive free energy difference of 4.96 and 4.79 kcal/mol for DHF and TMP binding, respectively. The A26T mutation, on the other hand, contributes a negative free energy difference of -1.90 and -1.26 kcal/mol for these cases. In the absence of epistasis, we expect the presence of the A26T mutation not to be able to alleviate the destabilization due to the W30G mutation. However, the actual difference is only 0.15 and -0.28 kcal/mol, i.e., the two mutations together are thermodynamically similar to the WT. This means that there is a large epistatic rescue of - 2.91 and -3.79 kcal/mol for the respective cases of DHF and TMP bound forms. We argue that this kind of epistasis is not dependent on the binder but is due to the conformational changes occurring directly on the protein chain. We display this outcome schematically in Figure 4a whereby both residues A26 and W30, while being located near the binding site but with side chains pointing away from the substrate, may interact constructively to lead to the observed positive sign epistasis (Figure 4d). Nevertheless, these changes do not contribute to the final fitness landscape in the presence of the inhibitor since similar effects are observed regardless of the small molecule hosted at the binding site. Therefore, the apparent epistasis which is the difference between the expected and the actual ΔΔ*G*_*competition*_ is less than 1 kcal/mol (Table 5). Conversely, for the double mutation involving I94L and A26T which do not display any direct or indirect cross interactions, there is no/negligible epistasis in individual binding of substrate/inhibitor (Table 4); they also lack any apparent epistasis (Table 5).

**Table 5.** Role of epistasis in competitive inhibition.^†^.

| Mutants | $\Delta\Delta G$<br>(kcal/mol) | $\Delta\Delta G^e$<br>(kcal/mol) | apparent epistasis<br>(kcal/mol) |
| --- | --- | --- | --- |
| **L28R-A26T** | -4.33 ± 0.37 | -2.84 ± 0.37 | **-1.49 ± 0.53** |
| **L28R-W30G** | -3.35 ± 0.40 | -2.05 ± 0.41 | **-1.30 ± 0.57** |
| W30G-A26T | 0.43 ± 0.20 | -0.45 ± 0.21 | 0.88 ± 0.27 |
| I94L-A26T | -0.18 ± 0.12 | -0.73 ± 0.38 | 0.55 ± 0.40 |

<sup>†</sup> The superscript e indicates the expected free energy difference if the individual mutations were additive.

**Figure 4.**
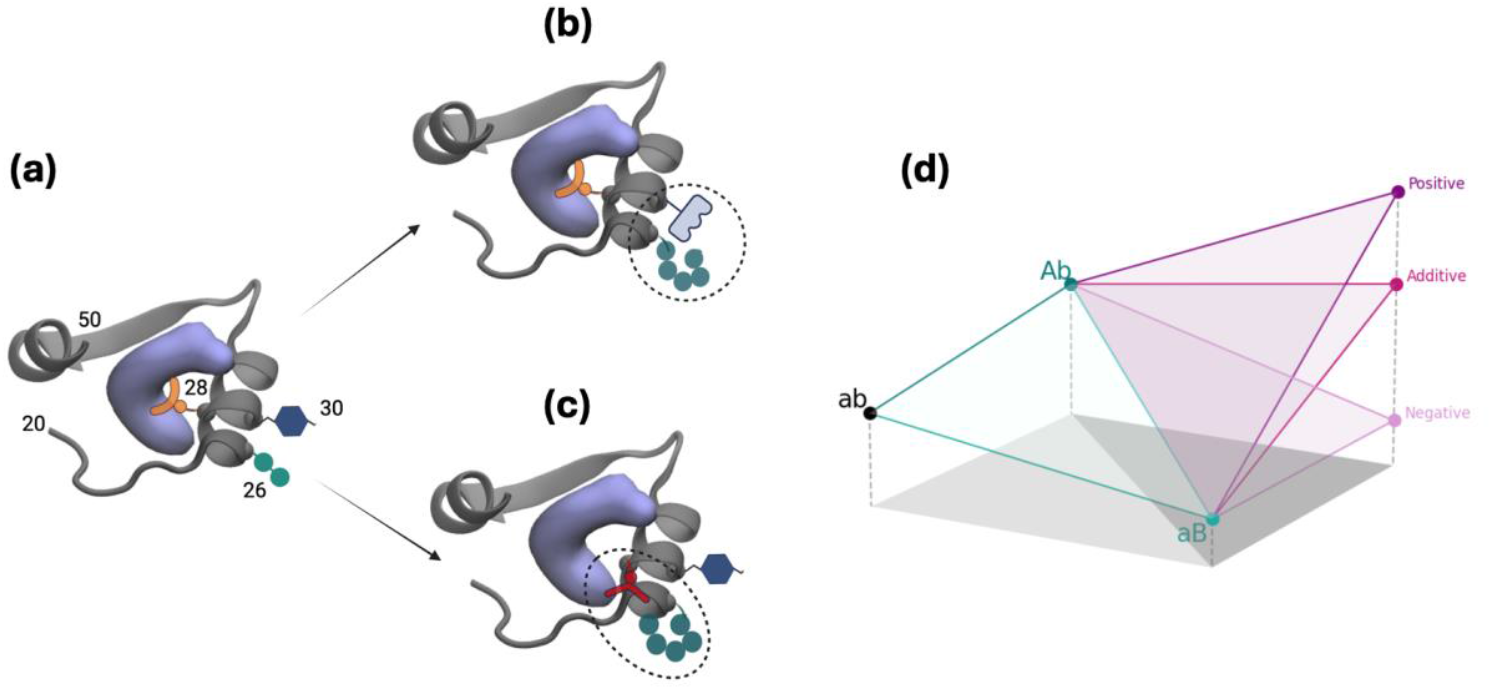
Schematic illustration of the effects of frequent mutations observed near the substrate binding site of DHFR for **(a)** WT, **(b)** double mutation at positions 26/30, **(c)** double mutations at positions 26/28. Direct or indirect interactions of the side chains with each other and/or the substrate may be responsible for epistasis. The protein structure (PDB code 1rx2) for residues 20-50 is shown in cartoon, the substrate DHF is shown as a volumetric purple blob and the positioning of the side chains of residues 26, 28 and 30 are illustrated by various geometric shapes. **(d)** Fitness assessment where larger values correspond to lower free energy difference; capital letters imply point mutations. Combining mutations can result in additivity (i.e., no epistasis) as exemplified by the I94L-A26T double mutation. An increase over the expected fitness, e.g. for the DHF-bound L28R-A26T system leads to positive epistasis, and negative epistasis is exemplified by TMP binding to the same system. (a)–(c) created with BioRender.com.

A different outcome is observed for both double mutants involving L28R that we have studied. These display smaller epistasis, on the order of -1 kcal/mol for the DHF bound forms and in the positive direction in the TMP bound forms. The case for DHF bound L28R-A26T is schematically illustrated in Figure 4b whereby the A26T mutation modifies the direct interaction of the R28 side chain with the ligand/inhibitor. While these changes are small for each binder, since the effect is in opposite directions for the ligand and inhibitor, the apparent epistasis is significant for these mutations. As the inhibitor is smaller than the substrate (molar mass 290 and 441 g/mol, respectively; also see Figure 1a and d) thereby establishing fewer contacts with the enzyme, we might in general expect such differentiated epistatic effects for mutations that directly affect the binding site.

### Some comments on achieving convergence

Accurately exploring the necessary conformational space for each alchemical state is vital to the success of these results.^41^ However, this becomes problematic when a significant conformational rearrangement is required to transition from the reference to the target binder–receptor complex. While extending simulation times and employing enhanced sampling strategies can help, their impact is varied and especially difficult when the free energy barriers between conformational states are high, or when adopting and maintaining the desired target conformation proves challenging. In Table 1, we have listed the number of simulations required to get converged free energy values for either the DHF-bound and TMP-bound forms. Our criterion for convergence was to have overlapping samples in all 32 windows for the forward and backward simulations when at least four FEP sets were merged in the BAR analyses. For some of these cases we have achieved such convergence in four sets, but we have had to increase the simulation sets up to 7 in many cases. Overlaps in windows are exemplified in Figure S1. The full set of outputs are also provided (please see Data Availability Statement for details).

Two of the mutations we have handled are charge changing; L28R and W30R. A recent systematic study has shown that charge changing mutations are particularly prone to convergence issues, while usual FEP protocols are adept in predicting consistent free energy costs of neutral side chain replacements.^15^ The significance of charge corrections due to electrostatic contributions to binding emerged due to Ewald summation requirements in periodic boundaries. We have used the protocol from the literature^26^ as described under the Methods section which greatly alleviated charge correction errors. This approach also provides an additional check on the convergence of the free energy differences if the cost of the Cl^-^ creation in water is converged, to a value of 83.0 ± 0.5 kcal/mol in this case. On the other hand, the calculated BAR errors are larger due to merging two simulations to predict the free energy change.

We had additional challenges when simulating the W30R mutation. We have shown in our previous work that the effect of the W30R change is due to the salt bridge established between the newly created R30 side chain and E139 on the β-sheet residing on the opposite side, further away from the binding region (compare Figure 5a and b).^10^ The separation between the two residues is thus greatly reduced with a salt bridge distance at a baseline value of 2 Å which samples values up to 7 Å during a 200 ns trajectory. It also leads to an overall shrinkage in the distance between the α-helix accommodating R30 and the β-sheet where E139 resides (Figure 5c). This situation is hardly reproducible during a FEP simulation due to the gradual onset of the interactions and the relatively short simulation times that may not allow for sampling all the possible interaction scenarios. We have therefore included a second set of simulations where we have carried out FEP simulations for the R30W mutations, starting from the salt-bridged orientations and used the negative value of the final free energy differences. Only when we have combined the two sets of simulations, have we been able to converge the overlaps in the *λ* windows along with converged ion creation value as described for L28R.

**Figure 5.**
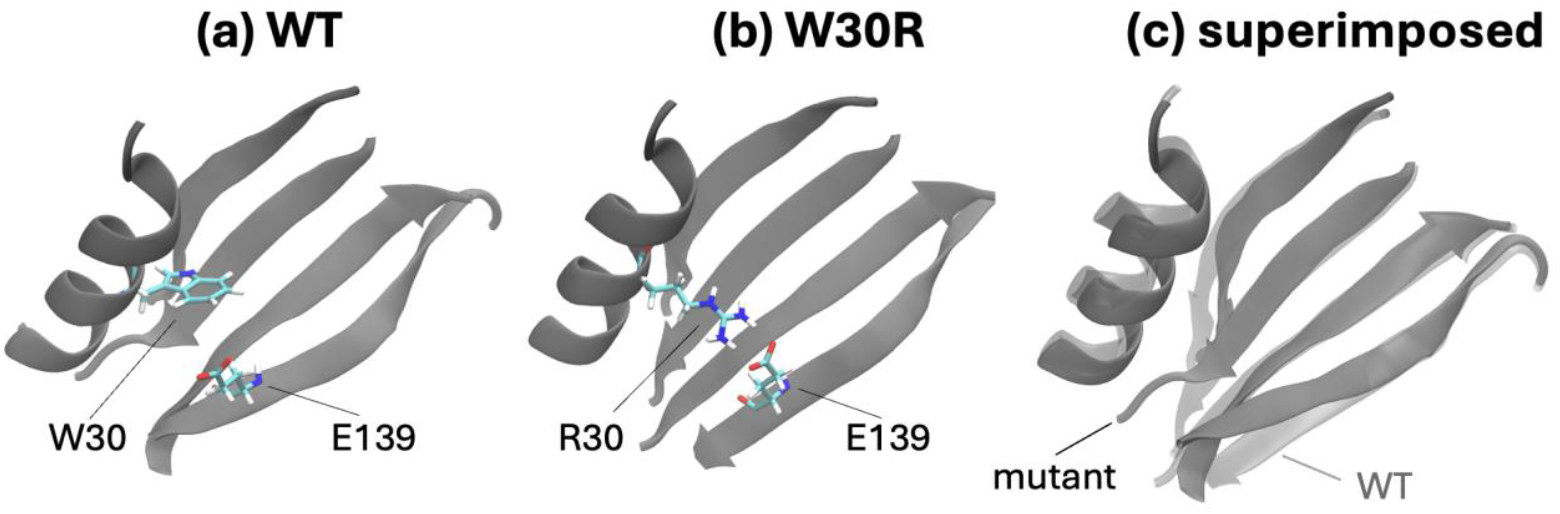
Relative positioning of residues 30 and 139 in the classical MD simulations of **(a)** WT and **(b)** W30R mutant. **(c)** The salt bridge established in the mutant ‘pulls’ the α-helix and β-sheet where these residues reside towards each other. The trajectories display a maximum 2 Å RMSD for the total protein.

Finally, the F153S mutation has proven to be another hard case. This residue is located near the C-terminus and is distal to the binding site. In fact, there is no apparent effect of the mutation that may by captured by comparing 200 ns long classical MD simulations for WT and F153S mutant via visual inspection of the trajectories. However, our previous work which followed large shifts in hydrogen bond occupancies have shown that this mutation has an allosteric effect on DHFR, interfering with the motions of the CD loop (Figure 1a) and revealing a cryptic site that is more than 30 Å away.^22^ As in the previous case of W30R, such long-range effects may not be captured by the gradual changes introduced into the side chains during the relatively short FEP simulations. Moreover, the overlaps in the first window of the reverse runs were very poor (Figure S1a), in the F153S simulations of the DHF-bound systems. Conversely, the S153F systems had well-behaved overlaps (Figure S1b), and we have reported the negative of the value predicted for these systems in Table 3. We did not have this problem for the TMP-bound system, hinting that the allosteric communication requires DHF.

For the double mutants, we have used the systems we have studied extensively in our earlier work as the base;^10, 21^ L28R for the L28R-A26T/W30G mutations; W30G for the W30G-A26T change and I94L for the I94L-A26T change. We have used the time points selected from the 200 ns MD simulations of the system with the single mutation to add the second mutation. We report the values in Table 3 as the sum of the costs of the two FEP simulations. However, we have also tried introducing both mutations at the same time for the I94L-A26T change.

We find that it is not straightforward to converge simultaneous mutations even for this case where both single mutants gave converged results in just four FEP simulations.

## Conclusions

The development of FEP-based methods addresses critical challenges in drug design, including accuracy, efficiency, and cost-effectiveness. By refining these methods, faster and more precise drug discovery processes may be achieved, ultimately improving patient outcomes and reducing development costs. FEP can model the interactions of drugs with mutated or polymorphic targets, aiding in the development of personalized therapeutics. For diseases with genetic variations such as cancer or antibiotic resistance, FEP holds immense potential in the design of tailored drugs. For high-throughput predictions, well-established computation protocols will prove essential.^15^

In this manuscript we develop a thermodynamic cycle which informs on the relative efficiency of an enzyme in the presence of a competitive inhibitor and relate it to the biochemical constants *K*_*m*_ and *K*_*i*_. We then demonstrate the applicability of this approach on 13 variants of *E. coli* DHFR that arise in bacterial colonies under the intense evolutionary pressure of TMP. Our thermodynamical cycle is based on FEP simulations carried out on substrate bound protein in the precatalytic conformation which has been argued to be the most significantly affected step by the mutations in enzymes. This is because, while mutations often shift population states of the enzyme, favoring catalytically competent conformations, these shifts do not necessarily extend to altering the pathways for product release, especially when product egress mechanisms rely on conserved structural features.^42^ This assumption holds well for our predictions, with 11 out of the 13 systems studied falling within the ±1 kcal/mol range (Figure 3), barring F153S and L28R-A26T, which are in the ±2 kcal/mol range. The measures of the overall deviations from experimental values, in terms of RMSD of the data (0.89), Pearson correlation coefficient (0.83) and AUC (0.82) are all favorable, considering the complexities of the subtle conformational changes and gradual ligand rearrangements. The largest discrepancy is for the F153S mutant whereby our simulations predict a negative ΔΔ*G*_*competition*_ value in favor of the substrate, while experimental data finds the difference negligible. We note that F153S is an allosteric resistance conferring mutation for which other mechanisms that come into play may not have been captured within the time frame of our simulations. Thus, the nuanced effects of mutations on protein behavior pose difficulties in predicting free energy changes, e.g., challenges emerge when compensating for charge changes, necessitating usage of additional runs, or simulation of the reverse mutation to cover the possible conformational range.

Our approach introduces a protein-centric view on the competitive inhibition mechanisms and allows us to interpret the effect of the mutations on the substrate-bound and the inhibitor-bound forms of the enzyme separately. Thus, substrate binding and inhibitor binding may be assessed individually, which is a big advantage for rational drug design purposes. In particular, similar free energy differences for both implies that the effect of the mutation does not directly involve the binding site. Moreover, epistasis may also be interpreted in a novel way and background mutation selection for high-throughput mutational scanning experiments may be suggested based on this additional structural information. With the currently available computational resources, carrying out a total mutational scan on the WT, followed by mutational scans in the background of mutations selected based on this detailed knowledge have become possible.^15^ This capability opens the way forward for interpreting epistatic effects on fitness landscapes.

## Data availability

All automated scripts for calculation setup and the outputs of the FEP simulations can be found at https://github.com/midstlab/cyclescript.

## ACKNOWLEDGMENTS

This work was financially supported by NIH project no. 5R01GM125748–07. The numerical calculations reported in this paper were partially performed at Scientific and Technological Council of Türkiye National Academic Network and Information Center (TUBITAK ULAKBIM), High Performance and Grid Computing Center (TRUBA resources).

## Notes

https://github.com/midstlab/cyclescript

### Competing Interest Statement

The authors have declared no competing interest.

