## Supplementary material for "A thermodynamic cycle to predict the competitive inhibition outcomes of an evolving enzyme": Figure S1

[†] Present address: Department of Chemistry & Biochemistry, University of Arizona
[‡] Present address: Department of Strategic Development, University of Twente

**Figure S1 |** Probability densities for the energy distributions in DHF bound, **(a)** F153S and **(b)** S153F runs.

**(a)**

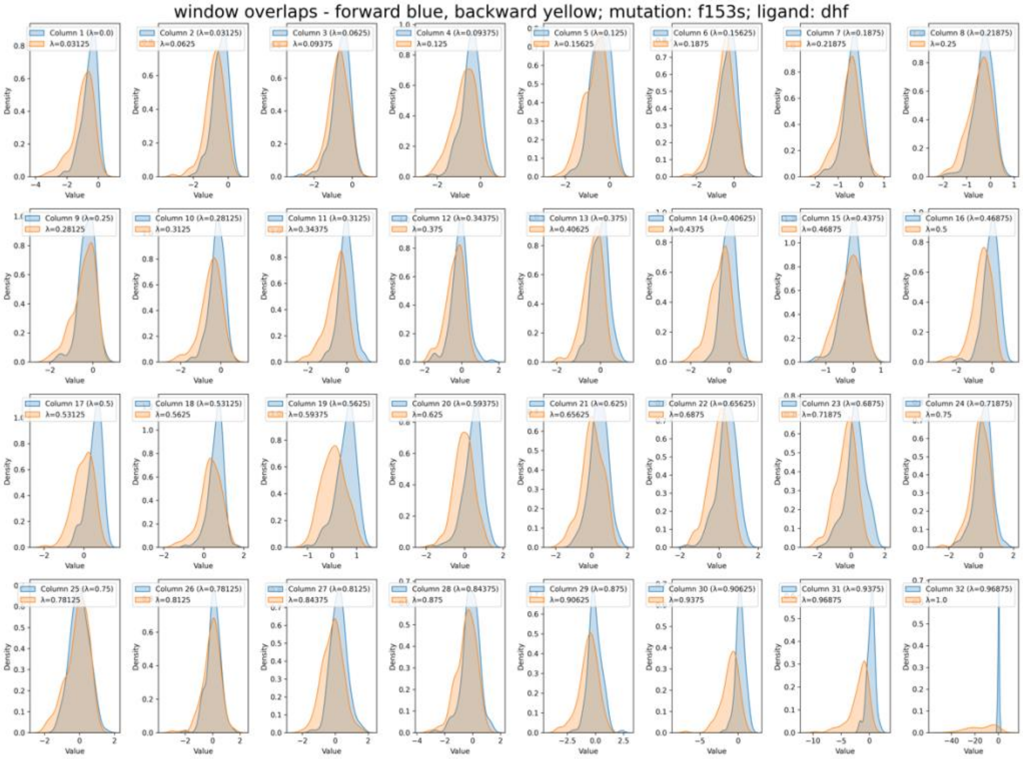


**(b)**

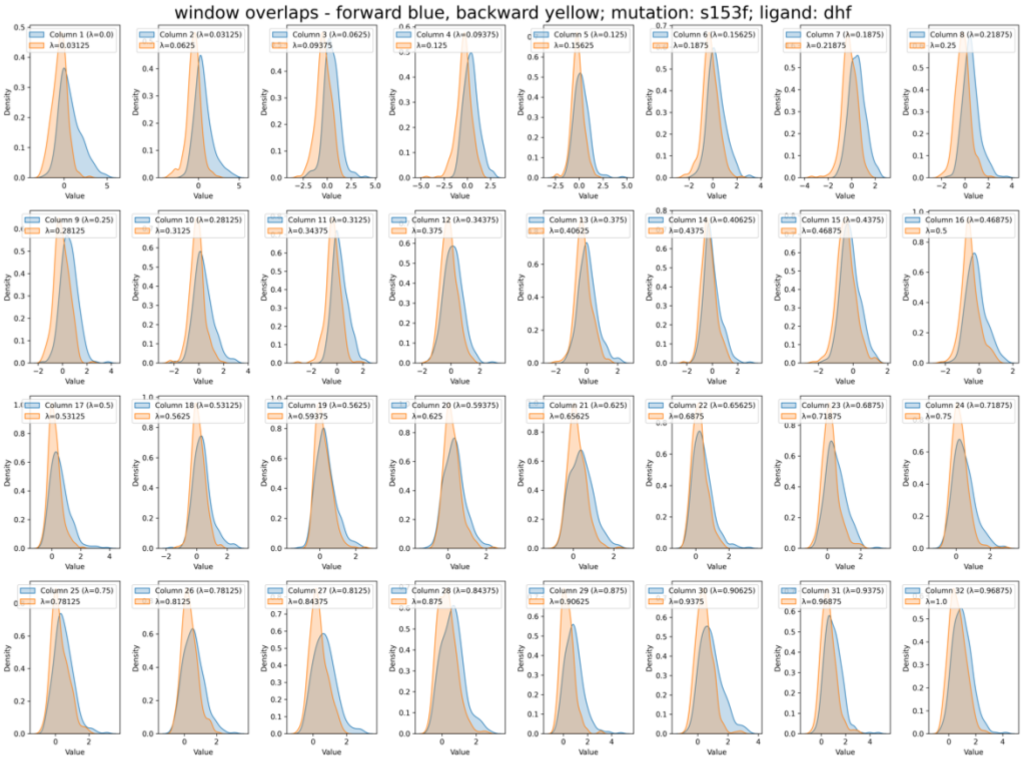
